# Opening the Pandora Box: DNA-barcoding evidence limitations of morphology to identify Spanish mosquitoes

**DOI:** 10.1101/354803

**Authors:** Delgado-Serra Sofía, Viader Miriam, Ruiz-Arrondo Ignacio, Miranda Miguel Ángel, Barceló Carlos, Bueno-Mari Rubén, Luis M. Hernández-Triana, Miquel Marga, Paredes-Esquivel Claudia

## Abstract

Cryptic speciation is frequent in the medically important mosquitoes. While most findings have been reported in tropical regions, it is an unexplored topic in countries where mosquito-borne diseases are not endemic, like Spain. The occurrence of recent outbreaks in Europe has increased the awareness on the native and invasive mosquito fauna present in the continent. Therefore, the central question of this study is whether the typological approach is sufficient to identify Spanish mosquitoes. To address this problem, we confronted the results of the morphological identification of 62 adult specimens collected from four different regions of Spain (La Rioja, Navarra, Castellón and the Island of Majorca) with the results obtained through DNA-barcoding. We conducted a comprehensive phylogenetic analysis of the *COI* gene region and compared this with the results of four species delimitation algorithms (ABGD initial partition, ABGD P=0.46%, bPTP and TCS). We report strong evidence for cryptic speciation in *Anopheles algeriensis* and *Aedes vexans* and reproductive isolation of the rock pool mosquito *Aedes mariae.* In addition, we report that the character present in the wings is not efficient to distinguish species *Culiseta annulata* from *Culiseta subochrea*, which distribution in the country may be different than previously described.

## 1. Introduction

Mosquitoes are responsible for the death of millions of people each year due to its role as vectors of some of the most devastating diseases in the world such as malaria, dengue and yellow fever. Although the majority of the cases are reported from tropical countries, several outbreaks have also taken place in the Northern hemisphere showing that the landscape of mosquito-borne diseases is changing in the world. In Italy, the rapidly-spreading invasive species *Aedes albopictus* have been implicated in outbreak of Chikungunya fever that affected 205 people in 2007 (Rezza *et al.*, 2007) and it was also identified as the vector of two Dengue fever cases in France (Gould *et al.*, 2010). Although it is unquestionable that invasive species represent a serious Public Health threat, the role of native mosquitoes should not be underestimated. Recent outbreaks of West Nile Virus (WNV) in Romania (Campbell *et al.*, 2001) and in the United States (Nash *et al.*, 2001), were attributed to local populations of *Culex* mosquitoes. Mosquito species identification has been traditionally carried out using dichotomous keys that describe the morphological characters of a particular life stage. Although this technique has proved valuable and is still widely used to distinguish many mosquito species, it has important limitations like the need of taxonomic experts to perform accurate identifications and the loss of morphological characters during collection and preservation of specimens. Furthermore, morphology-based taxonomy cannot differentiate indistinguishable members of species complexes, which frequently differ on their ecology, host preferences, behavioral patterns, therefore in their role as disease vectors, greatly impairing the design of effective control strategies (Paredes-Esquivel, Del Rio, *et al.*, 2009).

The field of mosquito taxonomy has being renewed with the development of new tools for accurate identification (Cywinska *et al.*, 2006). Molecular-based techniques are now widely used not only to identify sibling species but also to solve phylogenetic relationships (Bourke *et al.*, 2013). The *Cytochrome Oxidase I* gene region of the mitochondrial genome is the gold standard barcoding identification of species (Hebert *et al.*, 2003) and has proved valuable to distinguish many mosquito species (Cywinska *et al.*, 2006; Kumar *et al.*, 2007; Wang *et al.*, 2012; Ashfaq *et al.*, 2014). This approach is particularly important to establish species delimitations in unexplored groups (Puillandre *et al.*, 2012). Although, there have been reports that *COI* barcoding has failed in differencing some species of mosquitoes of *Anopheles* and *Culex* genus (Cywinska *et al.*, 2006; Kumar *et al.*, 2007; Wang *et al.*, 2012; Bourke *et al.*, 2013; Magdalena Laurito *et al.*, 2013), there is no doubt that DNA barcoding erects as a valuable alternative for mosquito identification in most areas of the world. However, the use DNA barcoding as a surveillance tool would require a database containing mosquito barcodes of correctly identified mosquitoes.

From the 3,552 formally recognized mosquito species in the world (Harbach, 2016), 100 have been recorded in Europe and this number might be even higher since six invasive *Aedes* species have recently colonized the continent (Versteirt *et al.*, 2015). Spain reports a similar increasing pattern, with 59 species reported at the beginning of this decade and 64 described in 2012 according to the latest checklist (R. Bueno Marí *et al.*, 2012), including competent vectors of human diseases. To the best of our knowledge, this would be the first comprehensive faunistic study that includes molecular-based techniques in Spain where identifications have mainly relied on the use of dichotomous keys. We are hoping to contribute to the knowledge of mosquito fauna and demonstrate that molecular based studies can uncover hidden diversity among well-known mosquito species.

## 2. Material and methods

### 2.1. Mosquito collection and morphological identification

The mosquito specimens were collected from 17 sites located in four different regions of Spain, including the Island of Majorca in the Balearic archipelago, Castellón, La Rioja and Navarra (Figure 1). The majority of the specimens (42) were collected from the island of Majorca, the largest at the Balearic archipelago, located in the Western region of the Mediterranean Basin. Additionally, for comparison purposes, three specimens from *Ae. mariae* were obtained from Peñiscola (Castellón, Spain), a coastal region with high abundance of salt marshes and rocky shores. Finally, 17 specimens from *Cs. annulata*, *Cs. subochrea* and *An. algeriensis* were collected from the wetlands of La Grajera and Hervías and in the River Iregua in La Rioja and Navarra, respectively. These sampling sites are located in the Ebro Valley, in the Northern region of Spain; with a transitional climate between the oceanic one in the North and a the semiarid found in the central Ebro Valley.

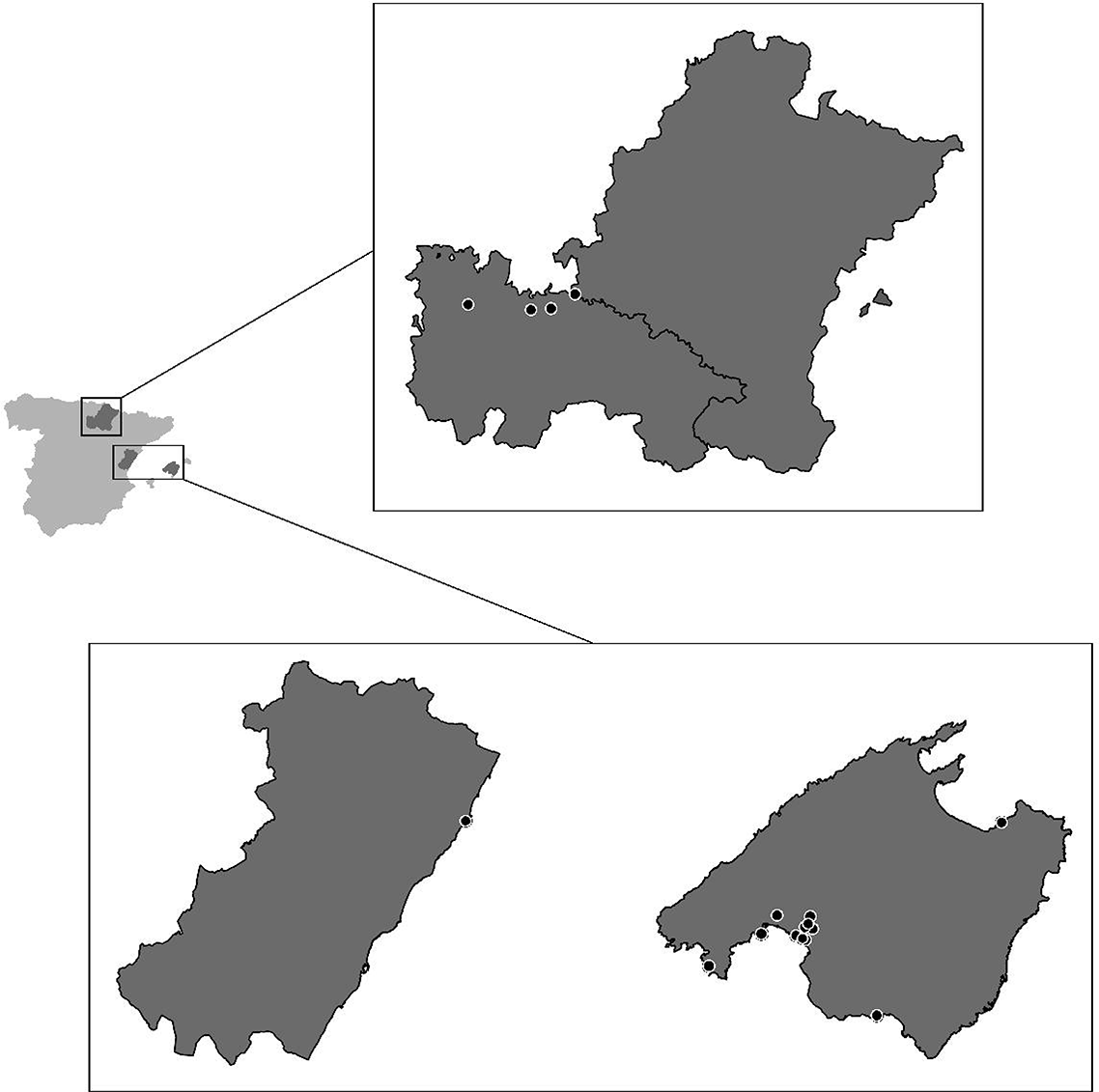

Adult mosquitoes were collected from outdoor using BG and CDC light traps containing CO_2_. The samplings were carried out weekly and each trap was set at sunset. On the following morning, collected specimens were transported *in vivo* to the laboratory and then frozen at −20°C for molecular studies. Larvae were also collected from breeding sites, transported to the laboratory and kept alive at 25°C until adults emerged. Female mosquitoes were identified to species level using keys by Becker et al. (2010). Scientific names of mosquitoes were verified using the Mosquito taxonomy inventory (Harbach, 2014).

### 2.2. DNA extraction, PCR amplification and sequencing

Genomic DNA was extracted from individual mosquitoes, usingDNeasy Blood & Tissue Kit (QIAGEN, GmbH, Hilden, Germany). To increase DNA yield we used 100μl elution buffer instead of 200μl in the final step. DNA concentration was measured with a NanoDrop 1000 spectrophotometer Thermo Scientific (Saveen Werner ApS, Denmark) and then DNA extractions were kept at -20°C (working solutions) and -80°C (Stock solutions). PCR was performed using the thermal cycler GeneAmp PCR System 2400 (Perkin Elmer) using primers C1-J-1718 forward 5 ‘- GGAGGATTTGGAAATTGATTAGTGCC-3’ and C1-N-2191 reverse 5’- CCCGGTAAAATTAAAATATAAACTTC-3’. Folmer primers (LCO1490 5’- GGTCAACAAATCATAAAGATATTGG-3’ and HCO2198 5’- TAAACTTCAGGGTGACCAAAAAATCA −3’) were also tested but failed to amplify all specimens. The 50 μl PCR reaction contained of 2 μl of DNA template, QIAGEN 1X reaction buffer (Tris-Cl,KCl) and (NH_4_)_2_SO_4_1.5–3.0 mM of MgCl_2_, 0.2 mM of dNTPs, 1 unit of QIAGEN DNA polymerase and 125 nM of each primer. The PCR reaction conditions were as follow: 95°C for 5 min followed by 92°C for 2 min and 15 s; 30 cycles of 92°C × 15 s, annealing temperature of 50°C × 15 s and an extension temperature of 72°C × 30 s, followed by a final extension at 72°C for 1 min. PCR products were stained with Pronasafe nucleic acid staining solution (Conda Laboratories) and observed in a 2% agarose - TBE buffer gel. Samples were purified using the QIAquick PCR Purification Kit (QIAGEN), according to manufacturer’s specifications, but purified DNA was eluted in 30 μl MilliQ water instead of 50 μl elution buffer and the template was sequenced in both directions in a ABI3730XL automatic sequencer (Macrogen Inc., Spain) with the same primers used in first amplification.

### 2.3. Sequence analysis

Sequences obtained were aligned using Clustal W and edited manually by using software BioEdit v7.0.8.0 (Hall, 1999). When point substitutions were detected, the reliability of the sequence was confirmed by looking at nucleotide peaks. All sequences were translated using the *Drosophila* genetic code in DnaSP version 5.10.1 (Rozas & Rozas, 1999) to ensure these were not pseudogenes using the mDNA. This program was also used to calculate haplotype diversity within species. The transition/transversion ratio and A+T content was calculated with Mega 6. For comparative purposes, we used BLASTn and sequences were retrieved from GenBank (https://www.ncbi.nlm.nih.gov/genbank/).

### 2.4. Phylogenetic analysis

The best evolutionary model to construct phylogenetic trees was selected according to their BIC score (Bayesian Information Criterion) obtained in Mega6 (Tamura *et al.*, 2007). With this program we also constructed the Maximum Likelihood (ML) tree and the robustness of the clustering was determined by the bootstrap analysis with 500 replicates. The Bayesian inferences were constructed with MRBAYES 3.2.6 (Ronquist & Huelsenbeck, 2003); using four Metropolis coupled Markov chain Monte Carlo (MCMC) chains were run for 500,000 generations (sampling every 10 generations) to allow adequate time for convergence. The analysis was finished when the standard deviation of split frequencies was less than 0.01. To assure we obtained the correct topology, analyses were run twice. All analyzes were performed with random starting trees. Branch support is indicated by the posterior probability values. Consensus trees were visualized with FigTree v1.4.3.

### 2.5. Species boundaries

We used the results obtained from the phylogenetic analysis as a first approach to distinguish species. Species were defined following the Phylogenetic species concept, that establishes that these are “irreducible cluster of organisms, within which there is a parental pattern of ancestry and descent, and which is diagnosably distinct from other such clusters” (Cracraft, 1987). The second approach we used to delimit species within our dataset was the Automatic Barcode Gap Discovery algorithm (ABGD), available at http://wwwabi.snv.jussieu.fr/public/abgd/. This method is independent of tree typology and split dataset into candidate species, based on the existence of barcode gap (interspecific divergence higher than intraspecific ones) and a prior intraspecific divergence (*p*) (Puillandre *et al.*, 2012). ABGD was carried out applying the K2P model, using default parameters, with the exception of the relative gap width, which was defined as 1.4. Results of a recursive and initial partition analysis were compared. The third approach to identify species was the Bayesian approach to Poisson Tree Processes model (bPTP) (Zhang *et al.*, 2013), which is based on the number of mutations in the branches of a phylogenetic tree. We run 100 000 trees, a thinning of 100 using a burn-in value of 0.1. A statistical parsimony networks were constructed using TCS (Clement *et al.*, 2000), which collapses DNA sequences into haplotypes and determines the frequency of those haplotypes within the dataset. Gaps were treated as 5^th^ state or missing data. Size of circles represents the number of sequences within a haplotype. Although this method has been mainly used to construct genealogies, it has been also used in diversity studies in other taxa. Molecularly Defined Operational Taxonomic Units (MOTUs) were defined according to the results obtained from the different species delimitation approaches. If these methods resulted in a different species composition, we defined MOTUs as those defined by the majority of such methods. We constructed a pairwise distance matrix using the Kimura 2-parameter model (K-2P) of substitutions using the aligned *COI* sequences in Mega 6 (Tamura *et al.*, 2007). We tested in the 10x threshold by Hebert et al (2004) to confirm morphological identification. This rule establishes that a new species is suspected if the average interspecific distance is at least 10 times higher than the average intraspecific divergence of the putative sister species.

## 3. Results

### 3.1. Sequence analysis

After alignment of the dataset, the matrix contained 406 bp long sequences. Neither insertion nor deletion events were observed in the dataset. All sequences were A+T rich, with an average content of 67.3% ranging 62.1% in *Coquillettidia richiardii* up to 69.2% in *Ae. mariae*, which is a similar value observed in other species of Culicidae (Paredes-Esquivel, Donnelly, *et al.*, 2009). From the 406 bp fragment, we found 147 parsimony informative sites and 10 singleton sites. Most substitutions were found at the third codon position, with a ratio of transitions/transversions of 1.13. Thirty-eight haplotypes were found in mosquitoes analyzed and their sequences were deposited in GenBank under accession numbers MH348258-MH348270 and MH370023-MH370049 (table 1). Haplotype diversity varied from 0 in *Ae. albopictus* up to 1 in *Ae. mariae* C, *Ur. unguiculata, Cq. richiardii, Ae. vexans* B and *An. algeriensis* B (Table 2). No evidence of pseudogenes was found as stop codons were not observed after translation, using the mitochondrial DNA genetic code for *Drosophila* sp.

**Table 1.**
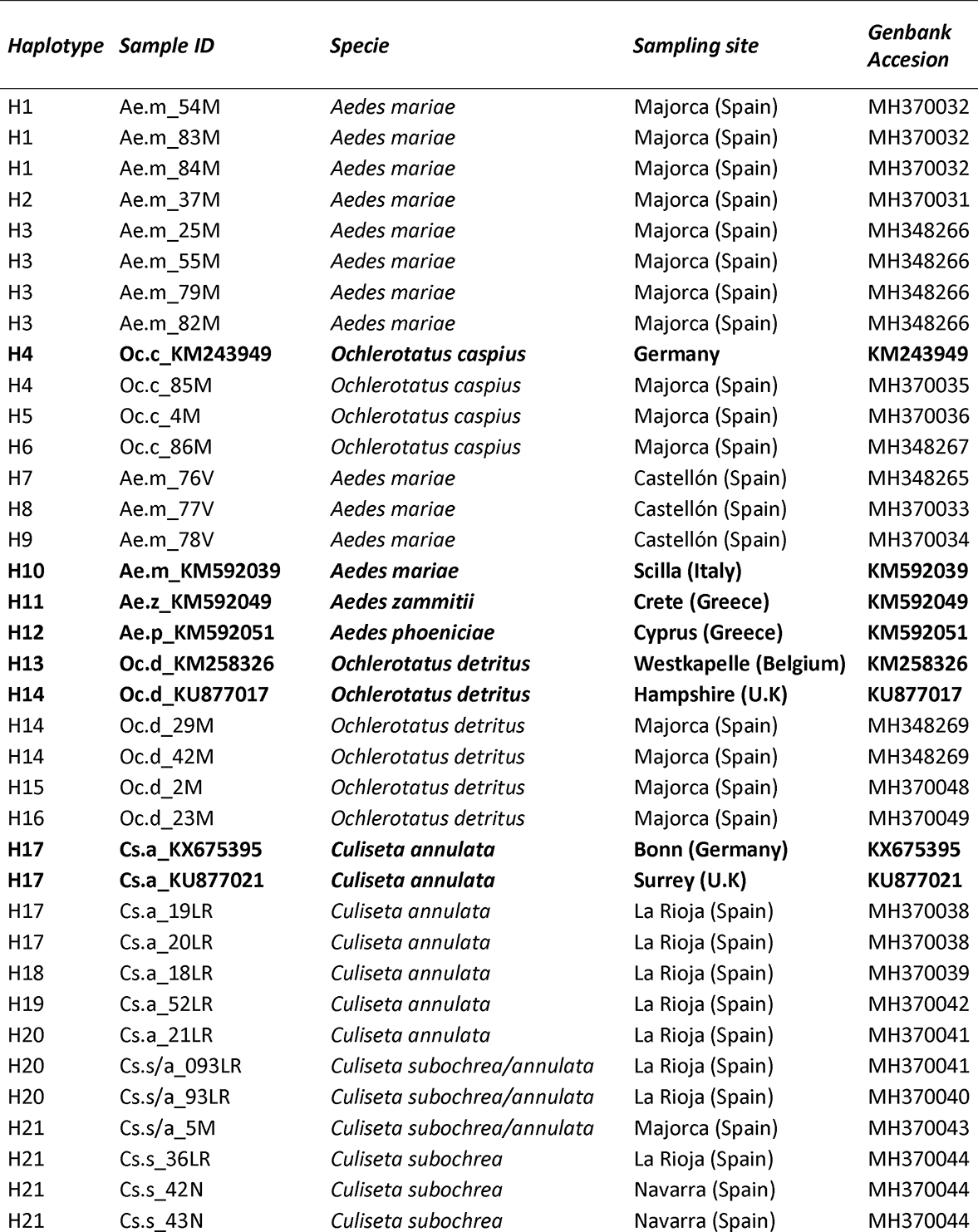
List of specimens included in this study

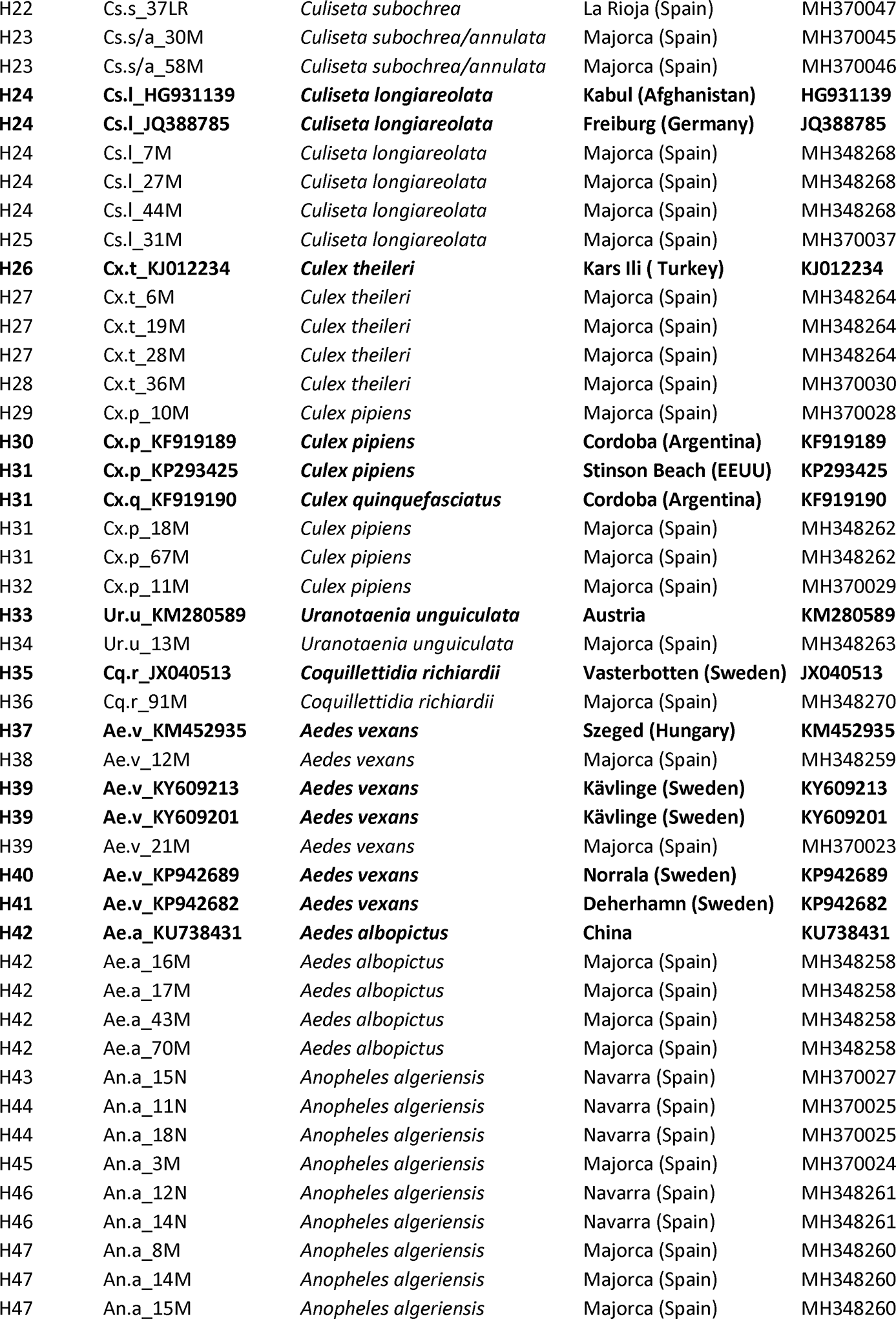

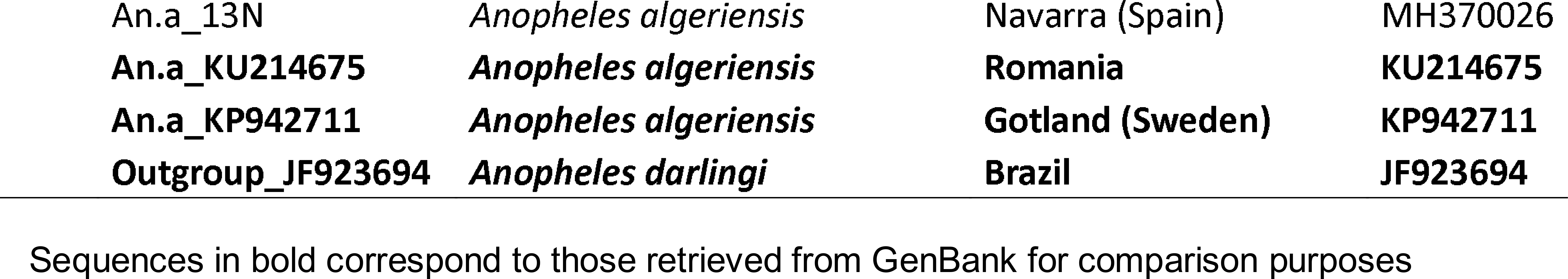

**Table 2.** Summary of genetic diversity parameters per mosquito species included in this study

| Species | N | SS | h | Hd (SD) | $\pi$ |
| --- | --- | --- | --- | --- | --- |
| *Aedes mariaae* M | 8 | 3 | 3 | 0.679 (0.122) | 0.00405 |
| *Ochlerotatus caspius* | 4 | 5 | 3 | 0.833 (0.222) | 0.00616 |
| *Aedes mariaae* C | 4 | 4 | 4 | 1.000 (0.177) | 0.00493 |
| *Ochlerotatus detritus* | 6 | 10 | 4 | 0.800 (0.172) | 0.00870 |
| *Culiseta annulata* | 9 | 3 | 4 | 0.750 (0.112) | 0.00233 |
| *Culiseta subochrea* | 7 | 2 | 3 | 0.667 (0.160) | 0.00188 |
| *Culiseta longiareolata* | 6 | 2 | 2 | 0.333 (0.215) | 0.00164 |
| *Culex theileri* | 5 | 2 | 3 | 0.700 (0.218) | 0.00197 |
| *Culex pipiens* | 7 | 12 | 4 | 0.714 (0.181) | 0.00891 |
| *Uranotaenia unguiculata* | 2 | 9 | 2 | 1.000 (0.500) | 0.02217 |
| *Coquillettidia richiardii* | 2 | 4 | 2 | 1.000 (0.500) | 0.00985 |
| *Aedes vexans* A | 5 | 4 | 3 | 0.700 (0.218) | 0.00443 |
| *Aedes vexans* B | 2 | 11 | 2 | 1.000 (0.500) | 0.02709 |
| *Aedes albopictus* | 5 | 0 | 1 | 0.000 (0.000) | 0.00000 |
| *Anopheles algeriensis* A | 10 | 4 | 6 | 0.889 (0.075) | 0.00421 |
| *Anopheles algeriensis* B | 2 | 11 | 2 | 1.000 (0.500) | 0.02709 |

N=number of samples, SS= Number of segregating sites, h=number of haplotypes, Hd=Haplotype diversity (Standard Deviation), $\pi$ = nucleotide diversity

Twenty-five sequences were retrieved from GenBank for comparative purposes (Table 2). Sequences belonged to the same species included in this study, with the exception of the Mariae Complex members *Ae. phoeniciae* and *Ae. zammitii* which were included to provide additional information on the *Ae. mariae* specimens from Majorca. A Kimura-2-parameter distance matrix was built for the entire dataset. The intraspecific divergence in the 406 bp fragment ranged 0.00-2.79 %, with an average 0.86%. On the other hand, distance among co-generic species ranged 2.13-15.74%. There was an overlap between the intraspecific and interspecific divergence (Table 3).

**Table 3.** Kimura 2-parameter of Molecular Operational Taxonomic Units identified in the phylogenetic analysis

|  | MOTUs | 1 | 2 | 3 | 4 | 5 | 6 | 7 | 8 | 9 | 10 | 11 | 12 | 13 | 14 | 15 | 16 | 17 | 18 |
| --- | --- | --- | --- | --- | --- | --- | --- | --- | --- | --- | --- | --- | --- | --- | --- | --- | --- | --- | --- |
| 1 | *Ae.mariae* M | **0,004** |  |  |  |  |  |  |  |  |  |  |  |  |  |  |  |  |  |
| 2 | *Oc.caspicus* | 0,02 | **0,006** |  |  |  |  |  |  |  |  |  |  |  |  |  |  |  |  |
| 3 | *Ae.mariae* C | 0,02 | 0,03 | **0,005** |  |  |  |  |  |  |  |  |  |  |  |  |  |  |  |
| 4 | *Ae.zammitii* | 0,04 | 0,04 | 0,03 | - |  |  |  |  |  |  |  |  |  |  |  |  |  |  |
| 5 | *Ae.phoeniciae* | 0,04 | 0,04 | 0,03 | 0,03 | - |  |  |  |  |  |  |  |  |  |  |  |  |  |
| 6 | *Oc.detritus* | 0,10 | 0,11 | 0,10 | 0,09 | 0,09 | **0,009** |  |  |  |  |  |  |  |  |  |  |  |  |
| 7 | *Cs.annulata* | 0,14 | 0,13 | 0,14 | 0,15 | 0,14 | 0,14 | **0,002** |  |  |  |  |  |  |  |  |  |  |  |
| 8 | *Cs.subochrea* | 0,13 | 0,12 | 0,13 | 0,13 | 0,13 | 0,14 | 0,05 | **0,002** |  |  |  |  |  |  |  |  |  |  |
| 9 | *Cs.longiareolata* | 0,12 | 0,12 | 0,12 | 0,13 | 0,12 | 0,14 | 0,13 | 0,14 | **0,002** |  |  |  |  |  |  |  |  |  |
| 10 | *Cx.theileri* | 0,12 | 0,13 | 0,12 | 0,13 | 0,13 | 0,14 | 0,15 | 0,14 | 0,12 | **0,002** |  |  |  |  |  |  |  |  |
| 11 | *Cx.pipiens* | 0,11 | 0,11 | 0,11 | 0,12 | 0,11 | 0,14 | 0,15 | 0,16 | 0,12 | 0,07 | **0,009** |  |  |  |  |  |  |  |
| 12 | *Ur.unguiculata* | 0,18 | 0,17 | 0,18 | 0,16 | 0,18 | 0,18 | 0,17 | 0,17 | 0,18 | 0,18 | 0,17 | **0,023** |  |  |  |  |  |  |
| 13 | *Cq.richiardii* | 0,21 | 0,20 | 0,21 | 0,22 | 0,22 | 0,22 | 0,21 | 0,21 | 0,20 | 0,19 | 0,19 | 0,23 | **0,010** |  |  |  |  |  |
| 14 | *Ae.vexans* A | 0,12 | 0,12 | 0,12 | 0,12 | 0,11 | 0,13 | 0,16 | 0,14 | 0,14 | 0,15 | 0,12 | 0,18 | 0,20 | **0,004** |  |  |  |  |
| 15 | *Ae.vexans* B | 0,12 | 0,12 | 0,12 | 0,13 | 0,12 | 0,14 | 0,18 | 0,15 | 0,15 | 0,15 | 0,14 | 0,19 | 0,21 | 0,06 | **0,028** |  |  |  |
| 16 | *Ae.albopictus* | 0,16 | 0,16 | 0,15 | 0,13 | 0,13 | 0,14 | 0,16 | 0,17 | 0,16 | 0,14 | 0,13 | 0,18 | 0,21 | 0,12 | 0,12 | **0,00** |  |  |
| 17 | *An.algeriensis* A | 0,17 | 0,17 | 0,16 | 0,15 | 0,15 | 0,21 | 0,21 | 0,21 | 0,17 | 0,17 | 0,17 | 0,21 | 0,24 | 0,18 | 0,20 | 0,18 | **0,004** |  |
| 18 | *An.algeriensis* B | 0,18 | 0,19 | 0,18 | 0,17 | 0,18 | 0,23 | 0,23 | 0,22 | 0,19 | 0,18 | 0,19 | 0,22 | 0,24 | 0,19 | 0,22 | 0,19 | 0,07 | **0,028** |

### 3.2. Phylogenetic analysis

Under the Bayesian Information criterion, the best evolutionary model obtained for our dataset was GTR+R. We used this model to the Bayesian analysis and substitution model by Tamura and Nei (Tamura & Nei, 1993) to construct the Maximum Likelihood phylogenetic tree. Both trees show clearly defined highly supported clusters (>87% and >96% in the Maximum likelihood and Bayesian trees, respectively) in all MOTUs identified (Figures 2 and 3). In both trees, members of the Mariae complex clustered in the same clade with *Oc.caspius. Aedes mariae* appears polyphyletic, with the Majorcan clade being more closely related to *Ochlerotatus caspius* than to *Ae. mariae* specimens from Continental Europe. Nevertheless, the relationship between these clusters was poorly supported by the bootstrap value. Specimens identified as *Ae.vexans* were separated in two highly supported clusters (>99%), herein *Ae. vexans* A, which included specimens from Majorca and sequences retrieved from GenBank from Sweden and Hungary. *Ae. vexans* B was formed by sequences from Sweden. These two groups have a 6% sequence divergence. Two monophyletic clades with high bootstrap support were also found in *An. algeriensis.* Those specimens from Majorca and La Rioja (*An. algeriensis* A) were clearly separated (bootstrap > 98%) from those sequences retrieved from Sweden and Romania (*An. algeriensis* B). These groups were separated by a 7% divergence (K2P). In addition all specimens from Majorca initially identified as *Cs. annulata* by field staff were placed in the *Cs. subochrea* clade (Figure 3). This resulted into a re-examination of the morphology of these specimens (see below). The remaining species *Aedes albopictus, Culiseta longiareolata, Culex theileri, Ochlerotatus detritus* and *Culex pipiens* formed highly supported monophyletic clades, although the last two showed some level of differentiation (Figure 2). Even when a single specimen of species *Uranotaenia unguiculata* and *Coquillettidia richiardii* were obtained, these were placed in a well-supported clade with those sequences from the same species retrieved from GenBank.

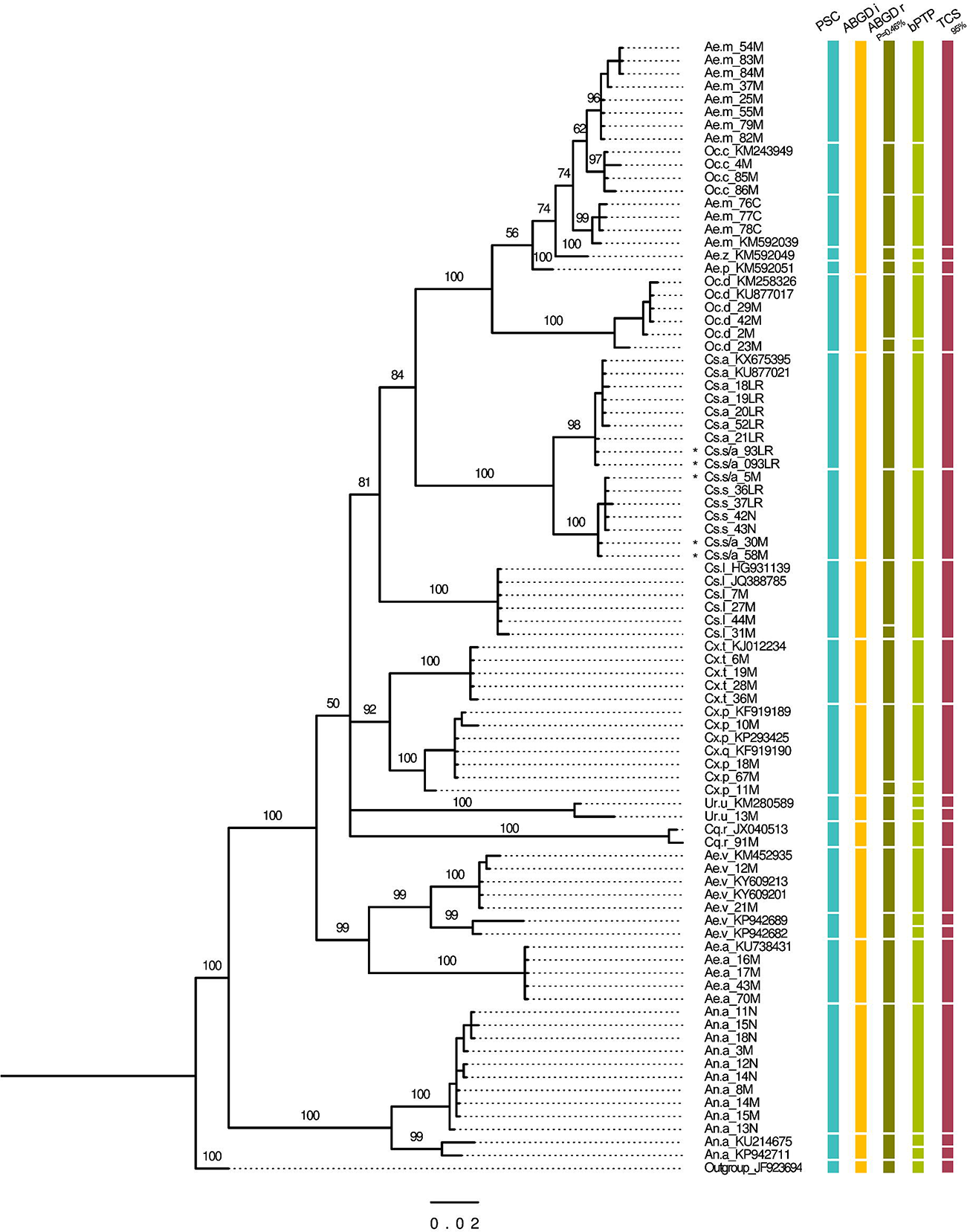

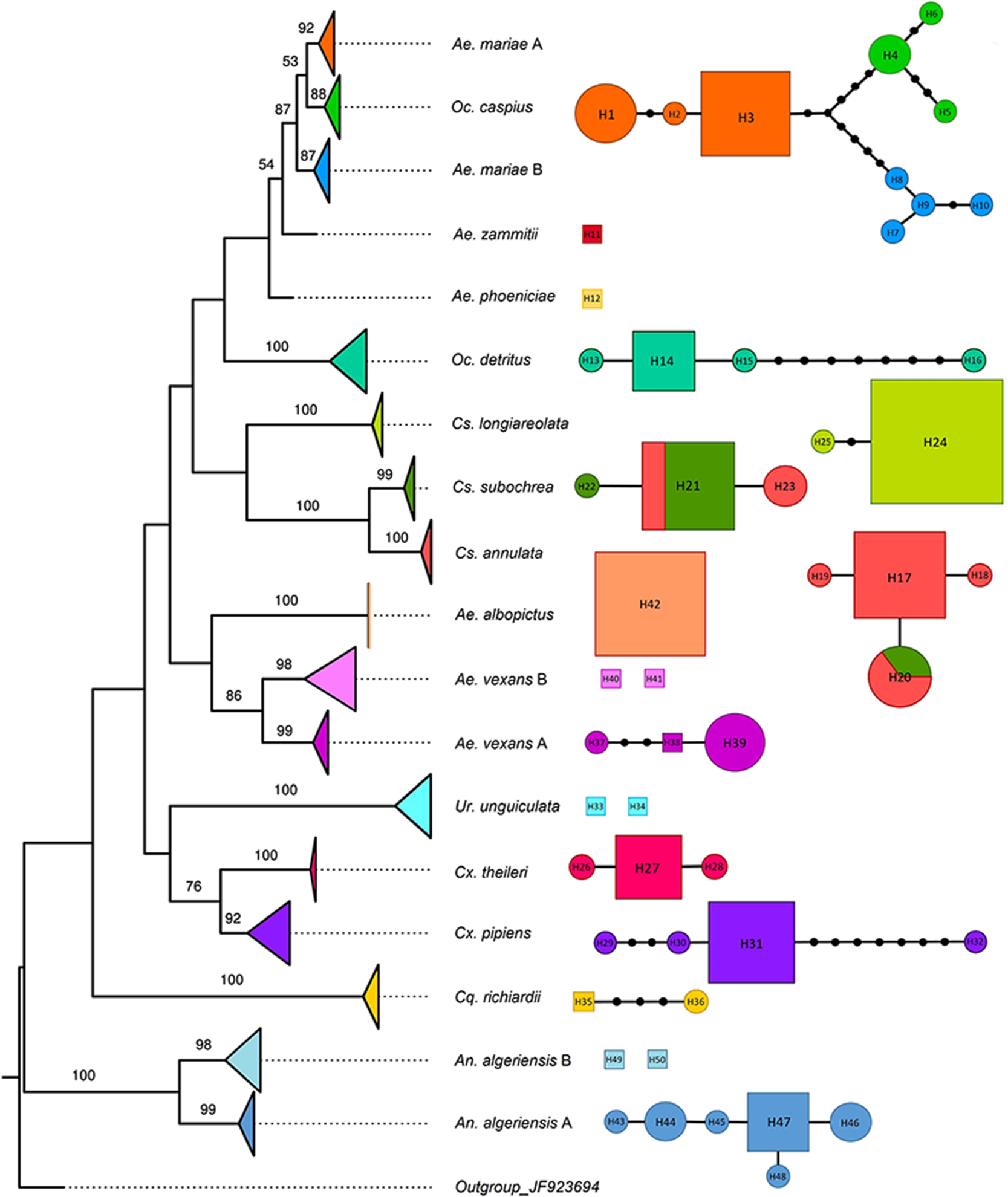

### 3.3. Species delimitation analysis

Using standard settings initial partition of the ABGD analysis resulted in 12 molecularly defined operational taxonomic units (MOTUs) for our 406 fragment of the *Cytochome Oxidase I* gene region. This first result clustered *Oc. caspius* and all members of the *Aedes mariae complex* (*Ae. mariae*, *Ae. zammitii* and *Ae. phoeniciae*) in a single MOTU and also failed to separate *Cs. annulata* from *Cs. subochrea*, while the remaining MOTUs corresponded to morphologically-identified species. On the other hand, this conservative approach split *An. algeriensis* in two species each. We also selected a recursive partition with a P value=4.64e-03, since this was the first to separate the three species of the Mariae Complex. However, when these were divided, the cluster containing *Ae. mariae* specimens from Majorca were also separated as a distinctive species (Figure 2). No significant differences were observed when using Jukes Cantor or Kimura K80. The bPTP analysis based on the Bayesian phylogenetic tree identified 23 MOTUs. The results of bPTP are mostly consistent with the ABGDr analysis; however, it is not as accurate as it separates some individuals as different MOTUs.

The last approach to separate species using the *COI* gene region was the genealogical haplotype network analysis using the statistical parsimony method implemented in TCS (Clement *et al.*, 2000). This method split our dataset in 19 networks separated by 11 mutational steps. The three species from the Mariae Complex were separated as different groups; however, *Oc. caspius* was included in the same network than *Ae. mariae.* The remaining specimens showed a clear agreement among haplotype networks and the morphological identification (Figure 3). There were clear disagreements in the number of species identified by each approach. However, most algorithms agreed in the separation of the MOTUs detailed in Figure 3. The Kimura 2-parameter showed that the majority of MOTUs identified by the species boundaries approaches could be identified using Hebert’s 10 X threshold (Table 3), with the exception of the members of the Mariae Complex and *Oc. caspius* from Majorca.

### 3.4. Agreement between molecular and morphologically-based identification

Using morphology-based keys we identified 13 mosquito species that belong to 7 different genera. Morphological identification using available keys were efficient to distinguish 11 of the species, with the exception of *Cs. annulata* and *Cs. subochrea.* Since the molecular identification resulted in conflictive arrangement of specimens, we re-examined the morphology of these and confirmed that all Balearic specimens showed morphological characters in the wings similar to *Cs. annulata* (“Cross veins r-m and m-cu aligned“), but abdominal scales like *Cs. subochrea* “Terga with indistinct pale basal bands formed by yellowish scales (not white), pale scales are also present among the dark scales in the apical half of the terga”. A somehow different situation was observed in La Rioja, with most specimens falling into the key’s description but some of them (Cs.a_93LR) with wings like *Cu. annulata* and terga like *Cs. subochrea* (Fig. 2). When characters located in the wings were ignored, the morphological identification was in agreement with the DNA-barcoding analysis.

## 4. Discussion

Establishing a genetic distance value in *COI* for species delimitation is a controversial topic that has been largely under scrutiny (Janzen *et al.*, 2017). Although a general value of 3% split for invertebrates species has been proposed, for mosquito identification the Barcoding community generally agrees with a Kimura-2-parameter value higher than 0.02 (>2%) to separate species (Kumar *et al.*, 2007; Ruiz-Lopez *et al.*, 2012). A low genetic divergence value (2,5%) has been used differentiate sibling species with different vectorial importance such as *Anopheles nuneztovari* from *An. dunhami*, a less important vector, in Colombia (Ruiz *et al.*, 2010). Closely related species within the *Anopheles albitarsis* group could also be differentiated using this value (Ruiz-Lopez *et al.*, 2012). Although this value may seem insufficient, a large-scale barcoding study conducted in butterflies, combining genomic and ecological data showed that a difference as shallow as 1 to 2% in the *COI* studied could be enough to differentiate species (Janzen *et al.*, 2017).

Using Bayesian and distance-based phylogenetic approaches and a combination of four additional species delimitation algorithms, we confirmed the identity of 11 out of 13 morphologically-identified mosquitoes (85%), demonstrating that DNA barcoding is an effective strategy to identify mosquitoes in Spain. These results are consistent with reports in other regions of the world, where congruence between these approaches ranged from 82% in Colombia (Rozo-Lopez & Mengual, 2015) up to more than 98% in most countries (Kumar *et al.*, 2007; Wang *et al.*, 2012; Ashfaq *et al.*, 2014; Chan *et al.*, 2014; Versteirt *et al.*, 2015). These differences may be related to the limitation of this technique in some taxa (M. Laurito *et al.*, 2017) or to errors in the initial morphological identification leading to incorrectly naming the sequences in the public databases used for sequence comparisons (Krzywinski & Besansky, 2003). We found disagreement between the barcoding and morphological approaches to separate *Cs. annulata* from *Cs. subochrea.* Furthermore, the sequence analysis also showed preliminary evidence of cryptic speciation in *An. algeriensis, Ae. vexans* and in *Ae. mariae s.s.* It is important to emphasize that the results obtained in this study are not definitive, as this only included one molecular marker. An inflation of the number of species due to the sole use of molecular methods is a risk to take into consideration (Harbach, 2004). Therefore, a combination of several markers is compulsory in order to arrive to firm conclusions on these potential cryptic species. Nevertheless, the DNA-barcoding approach has often served as a first step to identify cryptic species within Culicidae (Ruiz-Lopez *et al.*, 2012) and we hope that this will open a new window of research in this field in Spain, where identifications rely almost entirely in morphology. Results are discussed in depth below.

## 4.1. Preliminary evidence for cryptic speciation

All species delimitation approaches, including Hebert´s 10 x thresholds, agreed in the existence of at least two members within *An. algeriensis.* This is formed by two highly supported monophyletic clades separated by a high genetic divergence (K-2P = 6%). One of them, herein putative *An. algeriensis* A would be formed only by the Spanish specimens from Navarra and Majorca, while *An. algeriensis* B would be formed by specimens from Romania and Sweden and probably others not included in this study. *Anopheles algeriensis* is widely distributed in the Paleartic region, with Yemen as the easternmost limit (Al-Eryani *et al.*, 2016). This species is scarce in most areas where it has been reported (Krüger & Tannich, 2013), including the Balearic Islands. This scarcity led to the conclusion that *An. algeriensis* is a malaria vector of secondary importance (Becker *et al.*, 2010). However, this would not be the case of La Rioja, where it has been recorded as the third most abundant species (23,3% of all collected mosquitoes) in the wetland of La Grajera (Ruiz-Arrondo *et al.*, 2017). Furthermore, it has recently been found naturally infected with *Plasmodium vivax* in Yemen (Al-Eryani *et al.*, 2016), which should open new questions on their role in malaria transmission in endemic areas. In Spain, this species has been reported showing a “confusing” behavior, with variations in its host and resting preferences (R Bueno Marí *et al.*, 2011). Different patterns in mosquito behavior have been the first evidence of the possibility of cryptic speciation in other *Anopheles* mosquitoes (Paredes-Esquivel, Donnelly, *et al.*, 2009) and may be related to our findings. However these would not be surprising considering that half of the *Anopheles* species are part of species complexes (Harbach, 2004).

The use of nuclear and mitochondrial markers has recently allowed the detection of two different lineages within *Ae. vexans* (Lilja *et al.*, 2018). Our results are consistent with these findings, with all species delimitation analysis confirming the separation of at least two groups of *Ae. vexans* in the dataset. Mosquitoes collected from Majorca (Ae. *vexans* A) would belong to the authors’ Group 2, which has also been reported in Hungary and Sweden (Lilja *et al.*, 2018). *Ae. vexans* is a competent vector to more than 30 arboviruses, including West Nile virus, Zika virus and the dog heartworm *Dirofilaria inmitis* (Gendernalik *et al.*, 2017). In Spain it has been reported in 22 provinces, including the Balearic Islands (R. Bueno Marí *et al.*, 2012). *Ae. vexans* is a floodwater mosquito, with highly resistant eggs which can remain viable for several years. This species can travel long distances, which has direct relationship with its invasive capacity.

## 4.2. Incipient speciation in the Balearic rock pool mosquitoes?

The sea water contained in the rock pools of the Mediterranean coasts is habitat for mosquitoes of the *Aedes mariae* complex, which include three sibling species *Ae. mariae* s.s., *Ae. phoeniciae* and *Ae. zammitii* (Coluzzi & Sabatini, 1968). The phylogenetic analysis shows that *Ae. mariae* is not reciprocally monophyletic, with the Majorcan clade separated by >2% from the other members of the Mariae Complex, including the *Ae. mariae* specimens from mainland Spain and Italy. The Majorcan clade is also 2% distant to *Oc. caspius* and it appears more closely related to this species in the phylogenetic tree. However, the low bootstrap values obtained will not allow us to arrive to further conclusions. Contradictory results were obtained with the 10x threshold proposed by Hebert (2004) and the initial partition in ABGD. These approaches not only failed in separating the Majorcan clade, but also placed all members of the Mariae Complex plus *Oc. caspius* as a single species. These results are not consistent with the current knowledge of these species. The Mariae Complex has been subjected to a broad analysis, which included the morphological, genetic and reproduction studies of this group (Coluzzi & Sabatini, 1968; Mastrantonio *et al.*, 2016). There is no discussion on the species status of its members. However, it is known that they have a parapatric distribution with incomplete reproductive isolation (Mastrantonio *et al.*, 2016). Furthermore, including *Oc. caspius* and the Mariae Complex members in the same MOTU does not seem reasonable if we consider that these have a very distinctive ecology. While the former is a salt-marsh mosquito, the latter are rock pool mosquitoes with an extreme capacity to tolerate high salinity values. Further analysis using additional markers and running some experimental crosses in the laboratory would be necessary to establish the species status of the Majorcan *Ae. mariae* and its role as disease vector.

## 4.3. Current morphological keys cannot always distinguish Culiseta annulata from Cs. subochrea in Spain

Species *Cs. annulata* and *Cs. subochrea* are morphologically similar and can be differentiated by the presence of white (*Cs. annulata*) or yellow (*Cs. subochrea*) basal bands and the absence or presence of pale scales in the apical half of terga, respectively. In addition the alignment of veins r-m and m-c is effective to distinguish both species, being totally aligned in *Cs. annulata* and slightly separated in the case of *Cs. subochrea* (Becker *et al.*, 2010). The phylogenetic analysis and the species delimitation methods agreed in the clustering of two distinctive groups. As we obtained conflictive results with members of both species placed indistinctively in both clades (Figure 3), we re-checked the morphology and found that the character in the wings could not always separate both species, while the “the absence or presence of pale scales in the apical half of terga” was in clear agreement with DNA-barcoding identification. Conspicuous characters are frequently used by field workers as a careful examination of specimens can be time consuming. This may lead to erroneous delimitation of the distribution of these vectors. It is crucial to include other molecular markers to confirm these findings and re-design the morphological keys for these species in Spain. *Culiseta annulata* is a competent vector of the Tahyna virus vector (Bardos *et al.*, 1975) while we have found no reports on the vectorial capacity of *Cs. subochrea.* Therefore, its correct identification is crucial in case of outbreaks in the country. This would be the first record of *Cs. subochrea* in the Balearic Islands.

## 4.4. Disagreement among species delimitation methods to separate Spanish mosquitoes

We found disagreement among all species delimitation approaches used in this study. Methods such as ABGD and TCS allow the user to set the initial parameters, then results may be considered subjective. According to Puillandre *et al.* (2012) a P value = 0,01 is the closest to the species defined by authors when testing ABGD in four datasets. However, this value did not allow us to separate members of the *Aedes mariae* complex and *Oc. caspius* that appeared together in the same MOTU. On the other hand, bPTP, ABGD using a P value=4.64e-03 and the combination of the Phylogenetic Species Concept plus the 2% K2P approach were effective methods to separate these species. The fact that ABGD at initial partition and TCS were not able to separate members of the Mariae Complex, may be related to the incipient speciation of the group, which were more easily detected by the remaining species delimitation approaches. largely studied groups such as those from the Mariae complex is important as they may also be used as a standardization strategy to determine which species delimitation method is more appropriate for our dataset. In the case of *Oc. caspius* this would require a more careful analysis as this species seem close to *Ae. mariae* even when these vary in their habitats. Hybridization between these species cannot be ruled out.

A multilocus approach and a morphological revision is crucial to confirm the identification of these and other putative cryptic species and to confirm the validity of the morphological characters in *Culiseta annulata* and *Cs. subochrea.* As the identification approaches may likely disagree on the species composition, data on the ecological distribution the species should be incorporated as this may help clarify the species status as with *Oc. caspius*, which lives in salt-marsh mosquitoes, differing from the *Aedes mariae* complex species, which live in the rock pools of the Mediterranean coast.

## 4.5. Implications on disease transmission

Nine of the eleven morphologically-identified species have been recorded as vectors of human diseases. The remaining two are *Culiseta longiareolata* which rarely feeds on humans but it is considered a vector of veterinary importance and *Uranotaenia unguiculata* an autogenous species with no importance as a disease vector (Becker *et al.*, 2010). During the last years, there has been an increased awareness of mosquitoes and the diseases they transmit in Europe. Changes in climatic conditions together with globalization may result in active transmission of pathogens as it has already occurred in some areas. Their relative importance of such vectors depends on their feeding behavior, longevity, and other aspects that may vary in species complexes (Gendernalik *et al.*, 2017). Therefore, it is crucial to know which vector species are currently circulating in the continent. Identifying lineages within mosquito species has important epidemiological implications. For instance, the geographic limits of the lineages of the species *Culex annulirostris* has been associated to the transmission of Japanese Encephalitis Virus in Australasia (Hemmerter *et al.*, 2007). Cryptic speciation in mosquitoes is an underexplored topic in Europe and this study demonstrates that it may be present in many taxa. We hope this study will open the door to new studies in this topic.

## Acknowledgements

This work was funded by various sources: Mosquito collection was supported by the *Conselleria de Medi Ambient Agricultura y Pesca* of the *Govern of the Illes Balears*, DNA sequencing was possible thanks to *Ajuts a la recerca* for unfunded projects from the University of the Balearic Islands. We would also like to thank the *SOIB JOVE* program for funding the contract of the first author of the study and Mr. David Borràs for providing mosquito samples.

